# Exploring Differences in Functional Connectivity in Australian Rules Football Players : A Resting-State fMRI Study on the Default Mode Network

**DOI:** 10.1101/2024.09.15.613156

**Authors:** Mi Tran, Avanti Shrikumar, Sarah C Hellewell, Thomas Welton, Max Kirkby, Jerome J Maller, Paul Smith, Alan Pearce, Stuart M Grieve

## Abstract

The effects of non-concussive impacts in contact-sports such as in Australian rules football (ARF) are still largely unexplored. These impacts are often but not always lower in intensity, but occur more frequently than actual concussions. Since non-concussive impacts are often asymptomatic, their significance may be underestimated. Acute or subacute measurement of non-concussive injury is challenging as the pathological response and injury is poorly described. There is therefore a need for a greater understanding of the pathological consequences of exposure. Growing evidence indicates that resting-state functional connectivity (rs-fMRI) changes in the Default Mode Network (DMN) may be an important biomarker that is sensitive to characterize these impacts. In this work, we examined functional connectivity changes within the DMN of ARF players to evaluate its potential as an early biomarker for non-concussive impacts. Based on rs-fMRI, we compare the DMN of 47 sub-elite ARF players (mean age 21.5±2.7 years [SD], males 57%) and 42 age-matched healthy controls (mean age 23.2±2.3 years [SD], males 48%) using Independent Component Analysis (ICA) and Dual Regression. This approach permits an unbiased decomposition of brain activity into networks with principled handling of statistical error. An 83% increase in DMN connectivity (as measured by the Strictly Standardized Mean Difference on values derived from Dual Regression) was observed in ARF players in the left retrosplenial cingulate cortex compared to healthy controls (FDR-corrected p-value from dual regression = 0.03, 95% CI computed via bootstrapping was 58% to 116%). The AUC for distinguishing ARF players from controls was 0.80 (95% CI; [0.71, 0.89]), equating to a PPV of 78% and a NPV of 74%. These results are preliminary; future work could investigate robustness to different random initializations of ICA and validate the findings on an independent testing set, as well as investigate longitudinal changes in ARF players over the course of a playing season.

## 1. Introduction

Sports-related concussions (SRC), a form of mild traumatic brain injury (mTBI), are of particular health concern. Worldwide there are approximately 42 million people suffering from mTBI every year (1). In contact sports such as in the Australian Rules football, concussion occurs at a rate of six concussions per 1,000 hours of gameplay (2). A recent report published by the Australian Institute of Health and Welfare (AIHW) further underscores the severity of this issue. They revealed that there were 481 hospitalizations due to concussions between 2020-2021 (3). However, the vast majority of concussions are not hospitalized, suggesting that the true prevalence of concussive injuries is likely significantly higher than reported. The symptoms of SRCs vary depending on the impact. However, it has been shown that even injuries that appear mild can result in long-term complications, such as chronic post-concussion syndrome (4). To further emphasize the issue, repeated concussions may increase the risk of developing major depressive disorders and contribute to an increased incidence of neurodegenerative diseases later in life (5), (6).

Traditional methods of diagnosing sports-related concussion and determining return-to-play rely primarily on a combination of clinical evaluation and cognitive testing (7). These assessments typically utilize standardized tools, such as the Sport Concussion Assessment Tool (SCAT) (8). Conventional imaging modalities, such as computed tomography and structural magnetic resonance imaging (MRI), are employed when there is suspicion of significant structural damage (9). These methods are effective for detecting structural changes, but do not directly measure brain connectivity and function.

Functional magnetic resonance imaging (fMRI), especially resting-state functional magnetic resonance (rs-fMRI), has emerged as a powerful tool in detecting changes in brain activity (10), (11). Resting-state fMRI, where the subject is measured at rest, allows the study of neural networks while the brain is not engaged in any specific task (12). This provides insight into the intrinsic functional architecture of the brain. Resting state fMRI is based on measuring the blood-oxygen level dependent (BOLD) signal (13), a relative measure based on the ratio of oxygenation and deoxygenation of the protein hemoglobin. Increase in brain activity shifts the ratio towards oxygenated blood, which is detected by the MRI scanner (14).

There is some emerging evidence that the Default Mode Network (DMN) is altered following concussion (15), (16), (17). The DMN is predominantly activated when at rest or daydreaming - when the brain is not engaged in attention-demanding tasks (18). It is one of the most studied resting-state networks (RSN) (19). The Default Mode Network includes brain regions such as the posterior cingulate cortex (PCC), medial prefrontal cortex (MPFC), precuneus, and the parietal lobules (20), (18). Studies have shown that disruptions of the DMN are often observed in individuals with neurological disorders, including those who have suffered from mild traumatic brain injuries (21), (22). In the context of sports-related concussions, however, the impact on the DMN remains ambiguous. Research findings vary widely, likely due to differences in methodology, cohort characteristics, time course and concussion type. Some studies have reported an increase (21), (23) in DMN activity post-concussion, while others observed a decrease (24) (25). It is reported that increased DMN connectivity is linked with depressive disorders (26).

In contact-sports like ARF, there is continuous high-exposure to impacts that could result in concussion. These so-called non-concussive impacts may not meet the criteria for the diagnosis of clinical concussion. However, it has been hypothesized that brain damage may occur after repetitive exposure even in the absence of clinical concussion symptoms (27). In addition, little is known about the long-term effects of frequent exposure to non-concussive impacts.

Additionally, in the past decade, there has been a growing number of women participating in contact sports, e.g. in the ARF (28). In previous research by (29)), it has been shown that women suffer longer from cognitive impairment due to concussion compared to men. However, this was one of the few sports-related concussion studies involving women (30), (31). To account for sex differences in SRCs, a more targeted approach can be conducted in (for example) the decision-making in the return-to-play.

The aim of this study was to close the gap in our understanding of the impact of non-concussive events on brain functionality by assessing the changes in the Default Mode Network. Sub-elite female and male ARF players’ functional connectivity within their DMN were compared with age-matching healthy controls. It was hypothesized that even in the absence of clinical diagnosed concussions, we would still expect alterations in the DMN in the ARF players.

## 2. Methods

### 2.1. Participants

All participants provided written informed consent, with the study receiving institutional ethics approval prior to commencement from the Human Research Ethics Committee (HREC). Table I shows the demographics of participants and clinical features. 47 participants were recruited from a regional sub-elite ARF club, with teams in both the men’s and women’s first-grade competitions. ARF players were slightly younger than the healthy controls (student-t; p<0.05). There were no significant differences for weight and height when comparing ARF players with Controls grouped by sex. ARF players were scanned prior to the start of the ARF season. Healthy controls were excluded if they had history of mTBI and/or were engaged in any contact-sports.

**Table I.** Demographic and Clinical Features of the cohort.

| Clinical Features | ARF Players<br>(N=47) | Controls<br>(N=42) | Test-statistic | P-value |
| --- | --- | --- | --- | --- |
| **Sex, n (%)**<br>*Female*<br>*Male* | 20 (42.6)<br>27 (57.4) | 21 (52.4)<br>20 (47.6) | $\chi^2 = 0.281$ | p=0.596 |
| **Age years, mean (SD)**<br>*Female*<br>*Male* | 22.1 (3.03)<br>20.9 (2.23) | 23.4 (2.34)<br>22.9 (2.28) | t = 1.51<br>t = 3.03 | p=0.138<br>p=0.004 |
| **Weight kg, mean (SD)**<br>*Female*<br>*Male* | 65.9 (8.77)<br>78.5 (5.29) | 65.0 (13.0)<br>75.4 (13.3) | t = -0.275<br>t = -0.935 | p=0.785<br>p=0.361 |
| **Height meters, mean (SD)**<br>*Female*<br>*Male* | 1.67 (8.90)<br>1.82 (6.14) | 1.67 (7.92)<br>1.80 (6.60) | t = 0.0463<br>t = -1.09 | p=0.963<br>p=0.285 |

$\chi^2$ : Chi-squared Test | t: Student's-t

### 2.2. MRI data acquisition

rs-fMRI single-shot echo planar imaging (EPI) were obtained using a 3T GE SIGNA Architect MRI scanner (General Electric Healthcare, Milwaukee, Wisconsin, USA) at Epworth Medical Imaging (Epworth Hospital, Geelong, Australia) running DV26 software and using a 32-channel Nova head coil (Nova Medical Inc., Wilmington, Massachusetts, USA). During the rs-fMRI acquisition, participants were asked to relax and not to think of anything specific. The participants could keep their eyes open, keep their eyes closed, or fixate on a specified location. Rs-fMRI acquisition parameters were as follows: the acquisition consisted of 400 brain volumes over time with dimensions of 80 × 80 × 44. Slice thickness=3 mm, echo time (TE)=30 ms, repetition time (TR)=1.1 s, flip angle (FA)=70°, a field of view (FOV)=240 mm, voxel size=1 × 1 × 1 mm, multiband factor=3, and total scan time=440 s.

Additionally, a high-resolution MPRAGE PROMO T1-weighted image was acquired to complement the rs-fMRI data with the following parameters: dimensions 164 × 256 × 170, voxel size=1 mm isotropic, echo time (TE)= 3 ms, repetition time (TR)= 7.7 s, flip angle=8°, inversion time=900 ms, and total scan time=303s.

### 2.3. Image preprocessing

All MRI data were converted to Nifti file format using dcm2niix software (v1.0.20230411) (32). Resting-state functional MRI (rs-fMRI) was processed with FMRIB Software Library (v.6.0.7.6 http://www.fmrib.ox.ac.uk/fsl) (33). The first ten brain volumes were discarded to achieve steady state of signal, resulting in 390 remaining volumes for analysis. Preprocessing steps included brain extraction (BET; (34)) to remove non brainy tissue and subject motion correction (MCFLIRT; (35)) to align the time series with the middle volume as reference. In addition, the images were subject to temporal filtering with a high-pass filter (0.01 Hz) to remove slow frequency drifts in the signal, and were spatially smoothed (FWHM=2mm) to increase the signal-to-noise ratio. Large motion outliers were detected using the motion confounds file that was generated when implementing *fsl_motion_outliers*, and those brain volumes were removed. Structural T1-weighted images were first cropped to remove lower head and neck using FSL (robustfov) before skull stripping using FreeSurfer (mri_synthstrip; (36). The rs-fMRI was registered to its T1W structural image before linearly co-registering to the standard MNI152 brain space for group comparison using FLIRT (37). Nonlinear registration using FNIRT was not used at the time because it appeared to produce unrealistic warping of the brains.

### 2.4. Independent Component Analysis (ICA)

Independent Component Analysis (ICA) is a technique to separate the rs-fMRI signal into independent non-Gaussian signals (38). ICA was performed on the preprocessed resting-state fMRI data using FSL’s MELODIC (Multivariate Exploratory Linear Optimized Decomposition into Independent Components version 3.15 (39)). Multi-session temporal concatenation was applied on the whole cohort (controls and ARF players). This means that the data for every subject was concatenated after the last time point (14). The number of independent components (ICs) was set to a fixed number of 55, since this is approximately one seventh of the time series of our dataset (40). The Default Mode Network was identified by cross-correlating the group-level map at a specific threshold with a resting-state network template by (41)). All the components that had the highest correlation with the template were visually verified to extract the component that was most likely to be the DMN.

### 2.5. Dual Regression

To obtain subject-specific spatial and temporal maps for the DMN component, dual regression was employed (42). The idea of dual regression is to find the subject-specific maps that best describe each component in the group-level ICA map using linear regression (14). Dual regression consists of two stages. The first stage is to compute a subject-specific time series for each ICA component that describes how the component’s activity changes over the course of the subject’s fMRI time series. In this stage, the group-level ICA components are used as spatial regressors to explain the subject’s fMRI BOLD signal (14). The second stage of dual regression uses the normalized time series obtained from stage I as temporal regressors to obtain subject-specific DMN spatial maps for each component (42). An illustrative example of the Dual Regression framework is provided in **Figure 1**.

**Figure 1:**
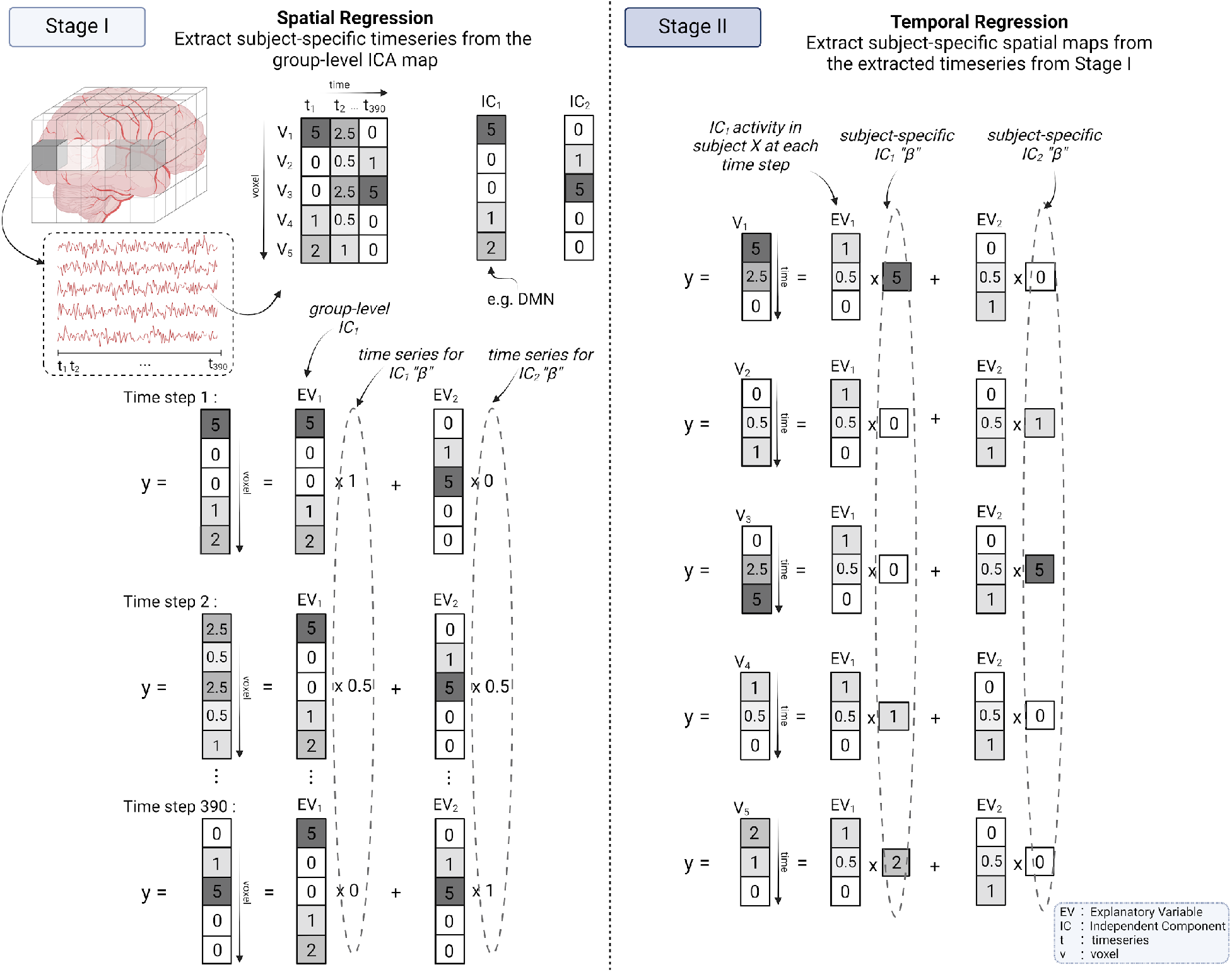
Dual Regression example for 5 voxels and 2 Independent Components. Stage I shows the Spatial Regression followed by Stage II Temporal Regression. The Beta values are fit in each stage. Note that in this example, the subject-specific components recovered in Stage II end up being identical to the group-level components.

For each ICA component that was chosen for further investigation, FSL’s randomise tool was then used to test the subject-specific spatial maps for group differences within their Default Mode Network connectivity (43). We used randomise with threshold-free cluster enhancement (TFCE) (44), which is a tool that computes spatial statistics in a way that does not require a fixed threshold to be set to define the clusters. TFCE computes a signal at each voxel that takes into account the signal of the surrounding voxels, the motivation being that the more similar the signal in the surrounding voxels, the more likely it is that a given voxel is part of a cluster. Specifically, for every possible threshold of signal strength that is less than or equal to the actual signal at a voxel, TFCE considers how many surrounding voxels meet that signal threshold, and then integrates a quantity derived from this value over all possible signal thresholds (44). Using the randomise tool, we performed permutation testing to determine the statistical significance of the differences between groups, with the number of permutations set to 5,000. Sex was included as a covariate in the permutation testing.

The randomise tool provides both raw p-values as well as p-values that are corrected for multiple comparisons across voxels using the Family Wise Error Rate (FWER). However, in our analysis, in addition to multiple comparisons across voxels, it was necessary to perform a multiple testing correction across all ICA components that were selected for further investigation, as well as to perform a multiple testing correction for the two possible directions of a group difference (both an increase and a decrease). As there were four ICA components chosen for further investigation, to perform this correction we applied a Benjamini-Hochberg FDR correction on the uncorrected p-values concatenated across all 8 spatial p-value maps (two p-value maps per ICA component, one p-value per voxel per ICA map).

## 3. Results

### 3.1. Independent Component Analysis

Out of the 55 components, we extracted 35 components that showed correlation above 0.2 with the ten resting-state network template (41). **Figure 2** shows the four components that mapped into the Default Mode Network (DMN).

**Figure 2.**
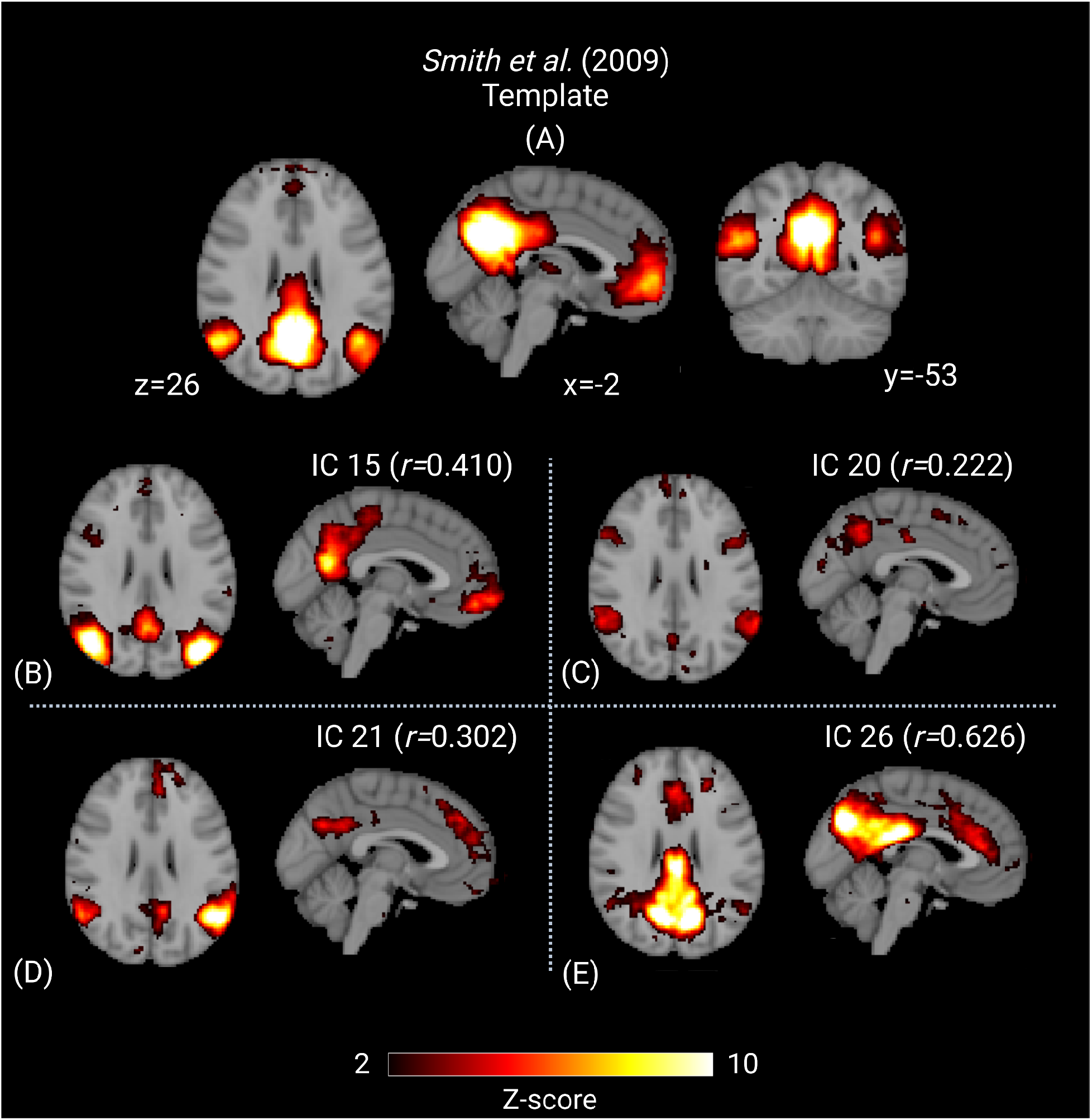
A: The Default Mode Network as defined by Smith et al., B-E: The four DMN components extracted from our group-level ICA map. The regions within the DMN shown include the posterior cingulate cortex (PCC), Precuneus, medial prefrontal cortex (MPC), and Left and right Lateral Parietal.

### 3.2. Comparison of Default Mode Network for Controls versus ARF Players

Permutation testing showed that there was significant difference (FDR=0.03) between the ARF players and healthy controls when comparing the functional connectivity within the Default Mode Network. The significance was found in ICA component 15 (see Fig. 2(B)). Statistically significant group differences between ARF players and healthy controls were not observed in the other three components that were mapped into the DMN. **Figure 3** shows the significant cluster within the DMN, with center MNI coordinates (−7, -38, 24). The significant cluster is localized in the left retrosplenial cingulate cortex, containing 8 voxels. This cluster was masked and the average beta values of all the voxels within that mask for every subject was extracted. ARF players show on average (mean±SD; 7.04±13.8) significantly higher functional connectivity within their DMN compared to healthy controls (mean±SD; -9.69±14.5). The corresponding histogram is shown in Figure 4. The ROC curve in Figure 5 demonstrates the model’s ability to differentiate between ARF Players and Healthy Controls based on average beta values, with an AUC of 0.80 (95% CI: 0.71-0.89), indicating good discrimination. The dashed line represents random performance (AUC = 0.5).

**Figure 3.**
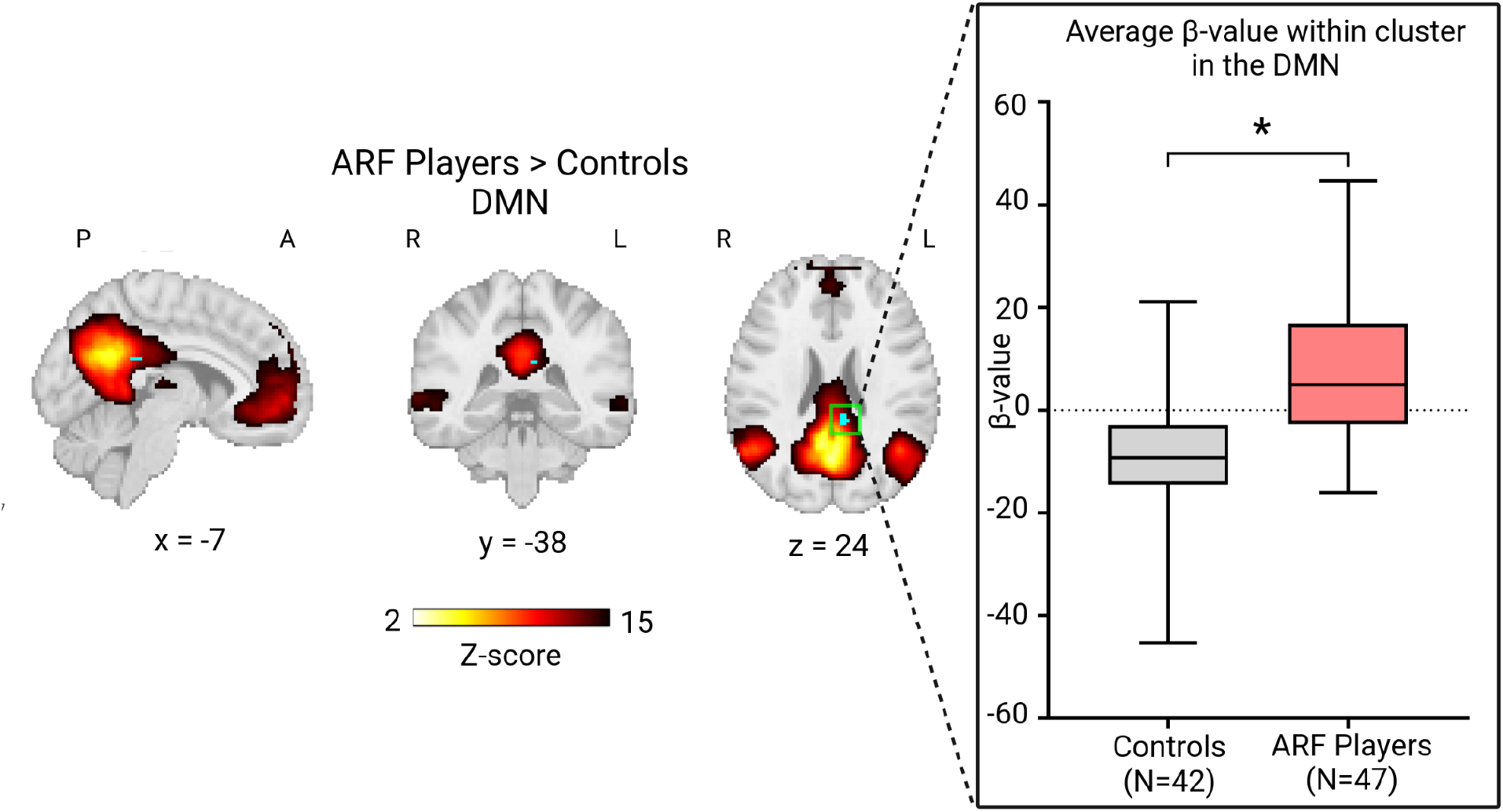
The significant cluster where ARF players show higher functional connectivity within the DMN compared to healthy controls. The boxplots show the Mean ± SD for each group. The p-value=0.03, FDR corrected.

**Figure 4.**
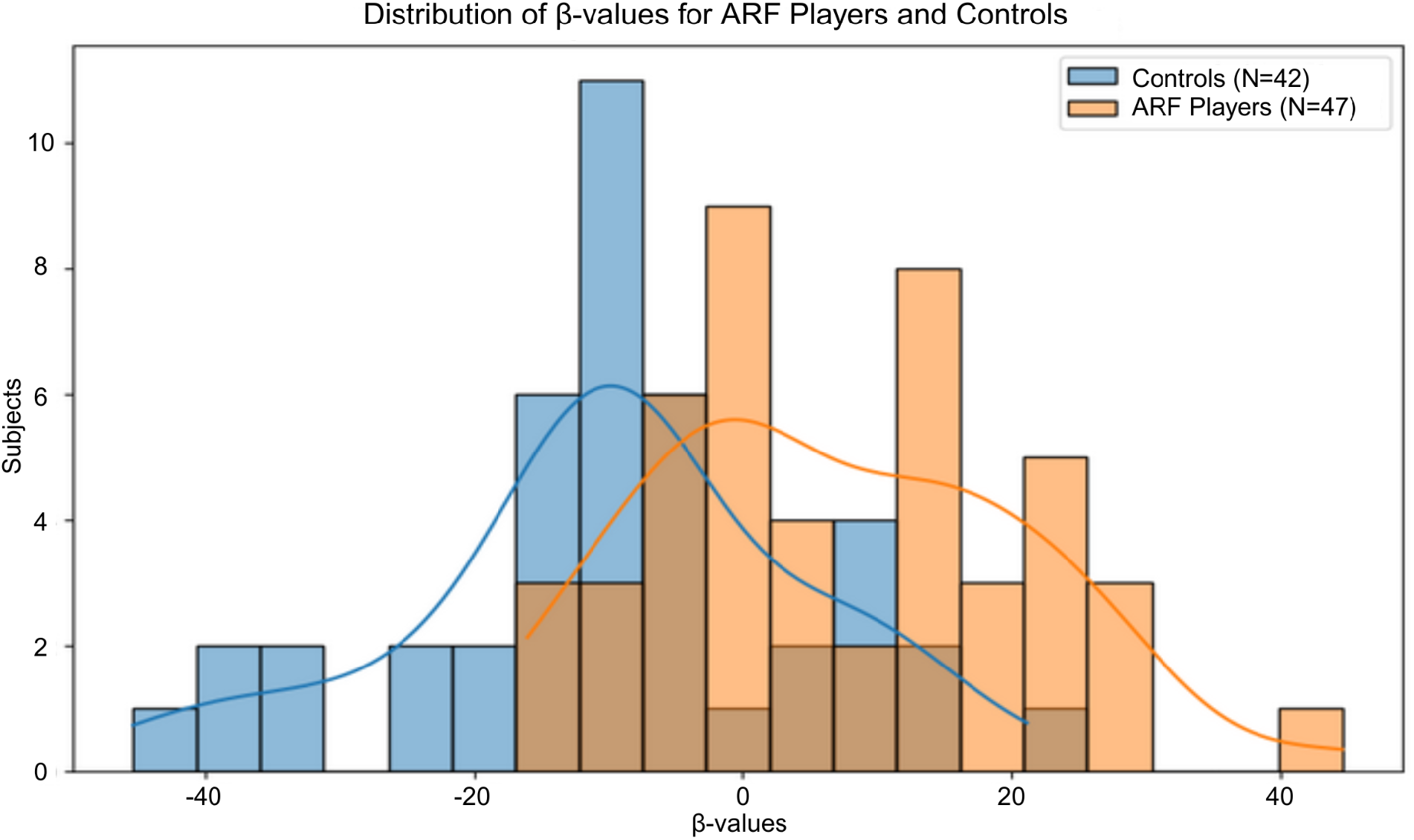
Histogram showing the beta value distribution of the ARF players and the healthy controls within the significant cluster.

**Figure 5.**
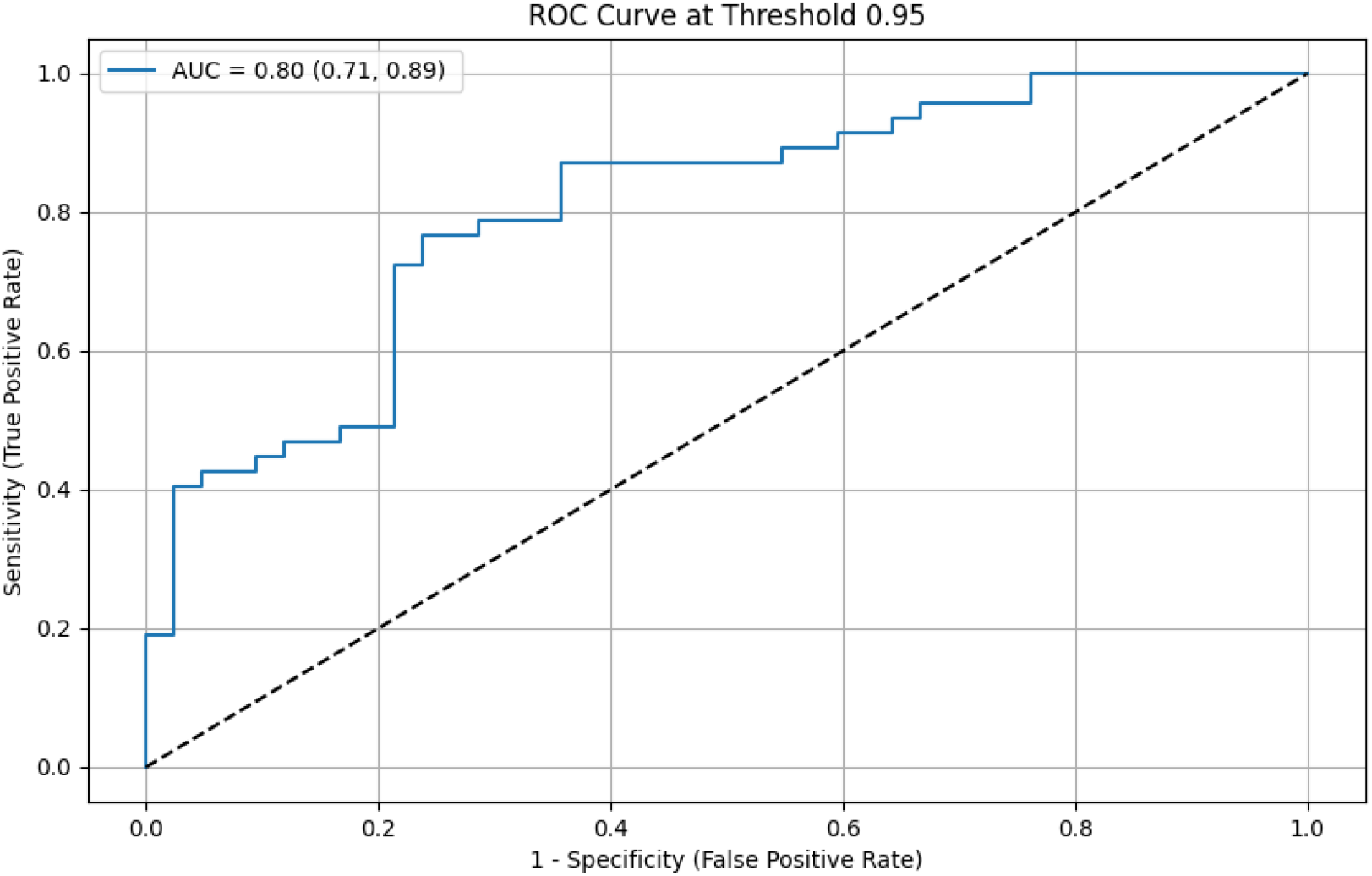
Illustration of the receiver operating characteristic (ROC) curve. The AUC value of the average beta values to differentiate between the ARF Players and Healthy Controls was 0.80 with a 95% confidence interval ranging from [0.71, 0.89]. The dashed line represents the performance at random (AUC=0.5). AUC: Area Under Curve | ROC: Receiver Operating Characteristic

## Discussion

In this study, we investigated differences in brain functionality between ARF players without a clinical diagnosis of concussion and healthy controls, using independent component analysis (ICA) and Dual Regression. Our main finding is that ARF players show higher activation in an ICA component mapping to the DMN compared to healthy controls. Our work aligns with the findings of a previous work by (45) that also used ICA to extract the DMN; in their longitudinal study comparing chronic traumatic brain injury (TBI) patients to healthy controls, they found increased functional connectivity within the DMN.

A study by (46) further supports our main finding. In their work, they showed that high school American football players have higher connectivity in their DMN prior to playing season as well as after playing season when compared to non-collision sports players. The authors identified a similar region in the left retro-splenial cingulate cortex as more active in players relative to controls The authors hypothesized that the higher connectivity both before and after the playing season was due to repetitive non-concussive impacts accumulated over years of participation in collision sports (46). In contrast to our study, the authors used a seed-based approach, which has known disadvantages compared to ICA as it does not derive the networks in a dataset-specific way and uses a predefined template to designate seeds within the DMN Regions of Interests (ROI). This process of defining seeds can be challenging, as small shifts in the seed can result in different findings (14). Additionally, seed-based analysis only considers one temporal signal at a time, while the brain networks are acting simultaneously and spatially overlap with each other (14). ICA is thus preferred as it can differentiate between these overlapping signals by considering all the network signals that make up the rs-fMRI signal (14). Another noteworthy difference between our study and the study by (46) is the size of the cohort. Our football players cohort is twice as large, and our healthy controls cohort is four times larger than their non-football players.

These results should be interpreted with caution, as they were not tested for robustness to the random seed of ICA, and were not validated on an independent testing set. Future work could address these shortcomings and investigate the impact of additional analysis techniques, such as employing denoising techniques such as ICA-FIX to remove the effects of unwanted variation in a more robust way.

In summary, our study showed that ARF players scanned before the playing season had a significantly higher Default Mode Network connectivity when compared to age-matched healthy controls, particularly in the left retro-splenial cingulate cortex.

